# Sumo-mediated recruitment allows timely function of the Yen1 nuclease in mitotic cells

**DOI:** 10.1101/2021.10.07.463465

**Authors:** Hugo Dorison, Ibtissam Talhaoui, Gerard Mazón

**Affiliations:** Université Paris-Saclay, UMR9019 CNRS, Gustave Roussy. 114, rue Édouard Vaillant, 94800 Villejuif, France; Inserm – Institut National de la Santé et de la Recherche Médicale, France; Hôpital Henri-Mondor, IMRB- U955 Inserm, 51 Av Maréchal De Lattre Tassigny, 94010 Créteil, France

**Keywords:** Homologous Recombination, Structure-selective endonucleases, Sumoylation

## Abstract

The modification of DNA damage response proteins with Sumo is an important mechanism to orchestrate a timely and orderly recruitment of repair factors to damaged sites. After replication stress and double-strand break formation a number of repair factors are Sumoylated and interact with other Sumoylated factors, including the nuclease Yen1. Yen1 plays a critical role to ensure genome stability and unperturbed chromosome segregation by removing covalently linked DNA intermediates that are formed by homologous recombination. Here we show how this important role of Yen1 is dependent on interactions mediated by non-covalent binding to Sumoylated partners. Mutations in the motifs that allow Sumo-mediated recruitment of Yen1 impair its ability to resolve DNA intermediates and result in increased genome instability and chromosome mis-segregation.

## Introduction

The DNA integrity of the genomes is constantly exposed to multiple challenges either from endogenous or exogenous sources of DNA damage. Cells have evolved multiple DNA repair pathways to ensure genome stability, Homologous Recombination (HR) being one of the most critical pathways to specifically counter the deleterious DNA double-strand breaks and other problems in the DNA integrity arising during replication. As the HR pathway operates, different DNA substrates and intermediates are formed that physically inter-connect distinct DNA molecules creating a joint-molecule (JM) intermediate. These intermediates are a threat to the successful segregation of chromosomes and are to be dismantled during mitosis by different specialized proteins acting in concert to prevent segregation defects and genome rearrangements. In yeast, the dissolution pathway mediated by the complex of Sgs1-Top3-Rmi1 (STR) ensures the disentanglement and release of double Holliday Junctions (dHJs) and two other helicases, Mph1 and Srs2, act early on the pathway preferentially over D-loop intermediates to reduce the number of JM intermediates and ensure the completion of the recombinational repair without crossing-over between the involved DNA templates. Opposed to these non-crossover (NCO) pathways, the nucleolytic processing of these JM intermediates can result in reciprocal crossovers (COs), with the risk of genome rearrangements and loss of heterozygosity (LOH) events (1, 2).

Given the risk for genome stability of a nucleolytic processing of HR intermediates, the different actors able to cleave these intermediates are strictly controlled and used as an option of last resort in DNA substrates not previously dismantled by the action of helicases (3, 4). Two major nucleases are involved in the nucleolytic processing of recombination intermediates in the yeast model, Mus81-Mms4 and Yen1 (5). The Mus81-Mms4 nuclease plays different roles at replication forks, and is gradually hyper-activated by Cdc5- and Cdc28/CDK1-dependent phosphorylation of Mms4 to peak its activity in late G2/M (6-8) where it associates with the Slx4-Dpb11 scaffold (9). Its broad substrate recognition enables Mus81-Mms4 to cleave 3’-flap containing DNA substrates and HJs, preferentially when they are still nicked or not completely ligated (10). Its hyper-activation in late G2 and its broad substrate specificity positions Mus81-Mms4 in a critical role to cleave both captured D-loop and early HJ intermediates, possibly targeting these intermediates that can’t complete full conversion to a dHJ and thus remain inaccessible to processing by the STR complex (3). As mitosis progresses, the Cdc14 phosphatase will trigger the reversal of the inhibitory Cdc28-mediated phosphorylation of Yen1 in turn allowing its nuclear localization and its proper substrate recognition (11, 12). This late activation of Yen1 at the anaphase entry ensures that all remaining recombination intermediates, especially those that escaped dissolution by STR or cleavage by Mus81-Mms4, are resolved before mitotic exit (11, 12). To ensure the clearing of Yen1 nuclease from the nucleus in the subsequent S-phase, and prevent off-targeted activity directed to 5’-flap containing DNA intermediates, Yen1 is additionally controlled by a Sumo-targeted degradation mediated by the Slx5-Slx8 ubiquitin ligase, further limiting the potential of crossover formation (13).

Protein covalent modification with the small ubiquitin-like modifier (Sumo) (14) is an important mechanism to fine tune DNA-mediated transactions during the DNA damage and repair responses (15-18). In *Saccharomyces cerevisiae* Sumoylation occurs in a multi-step reaction involving the E1 Aos1-Uba2 activating enzyme dimer, the E2 conjugating enzyme Ubc9, and three possible E3 ligases (Siz1, Siz2 and Mms21), with some redundancy of Siz1 and Siz2 for its substrates (19-21). Several players of the HR pathway, besides the nuclease Yen1, are also found among the Sumoylated DNA repair targets, including Rad52, PCNA, RPA and Sgs1 (17, 22-26). Sumoylation is able to influence biological processes in different ways. Proteins can be mono-Sumoylated, multi-Sumoylated or poly-Sumoylated, and the modification will re-design the protein surfaces allowing changes in protein activity, or in its way it can interact with other proteins. One of the best-described effects of protein Sumoylation is the enabling of interaction with other protein partners in a bait-to-prey fashion using Sumo as the moiety that is recognized by a specific domain in the partner protein, called a <u>Sumo I</u>nteracting <u>M</u>otif (SIM). These motifs are found throughout species and according to its amino acid composition can be classified in several families of consensus sequences (27). Most SIMs can be defined as a core stretch of four amino acids with a majority of hydrophobic residues (typically rich in V/I/L). This hydrophobic core fits into the hydrophobic groove on the Sumo surface and is often flanked by a stretch of 3-4 acidic or polar residues in the SIM sequence that interact with basic residues on the surface of Sumo (28-31). SIM types showing the flanking stretch of acidic residues present thus a similar architecture to that of ubiquitin interacting motifs (UIM) that also show a key stretch of polar residues flanking the hydrophobic core (32, 33). SIM motifs are able to interact with mono- or poly-Sumoylated proteins and can be usually present in tandem dispositions, probably helping interact with multiple Sumoylated lysines or with a poly-Sumoylated lysine in the interacting protein (28, 34, 35). Interactions by Sumo-SIM partnerships are extremely labile and can be easily induced and curbed down by altering the Sumoylation status of the involved proteins. This flexibility allows a quick building of protein complexes in response to changing stress conditions in the cell (17, 35). The build up of these protein complexes by Sumo-mediated recruitment *via* SIMs is generally associated to the actual Sumoylation of the two involved proteins (36, 37). Then, it was thus not surprising to identify in Yen1, which is Sumoylated (13), several putative Sumo interacting motifs. In the present study we define two functional SIMs in the Yen1 C-terminal region that play important roles in the nuclease sub-nuclear localization and its function alleviating the persistence of chromosome-segregation challenging JMs throughout the end of mitosis.

## Results

### Yen1 contains two functional Sumo-binding motifs (SIM) in the C-terminal region

As stressed before, Sumoylated proteins are able to transiently interact with a variable strength to other proteins containing SIMs. These motifs consist in a stretch of amino-acids with a core of aliphatic residues, often flanked by three or more amino-acids with a negative charge or susceptible to become phosphorylated (27). We inspected the Yen1 sequence through available algorithms (27) to detect SIM motifs and we identified several interesting hits in the primary sequence of Yen1 (Figure 1A). To validate the presence of such motifs in Yen1 we performed a two-hybrid analysis with Smt3 as bait and either wild type full-length or truncated versions of Yen1 as a prey (Figure 1B). The ability to interact with Sumo in the two-hybrid assay was only retained by the C-terminal part of Yen1 (amino-acids 354 to 759) and mutations in two putative SIMs present in that half of the protein completely abolished the interaction. A mutation in the first SIM, a type r motif (27) with a core at amino-acids 636 to 642 and a flanking stretch of acidic residues was indeed sufficient to almost completely impair the interaction to Smt3 in the two-hybrid assay (Figure 1B). Interestingly, this motif is highly conserved across Yen1 in other fungi and is very reminiscent of SIMs found in the Slx5, Rad18 and Elg1 proteins (36, 38, 39) (Figure S1).

**Figure 1.**
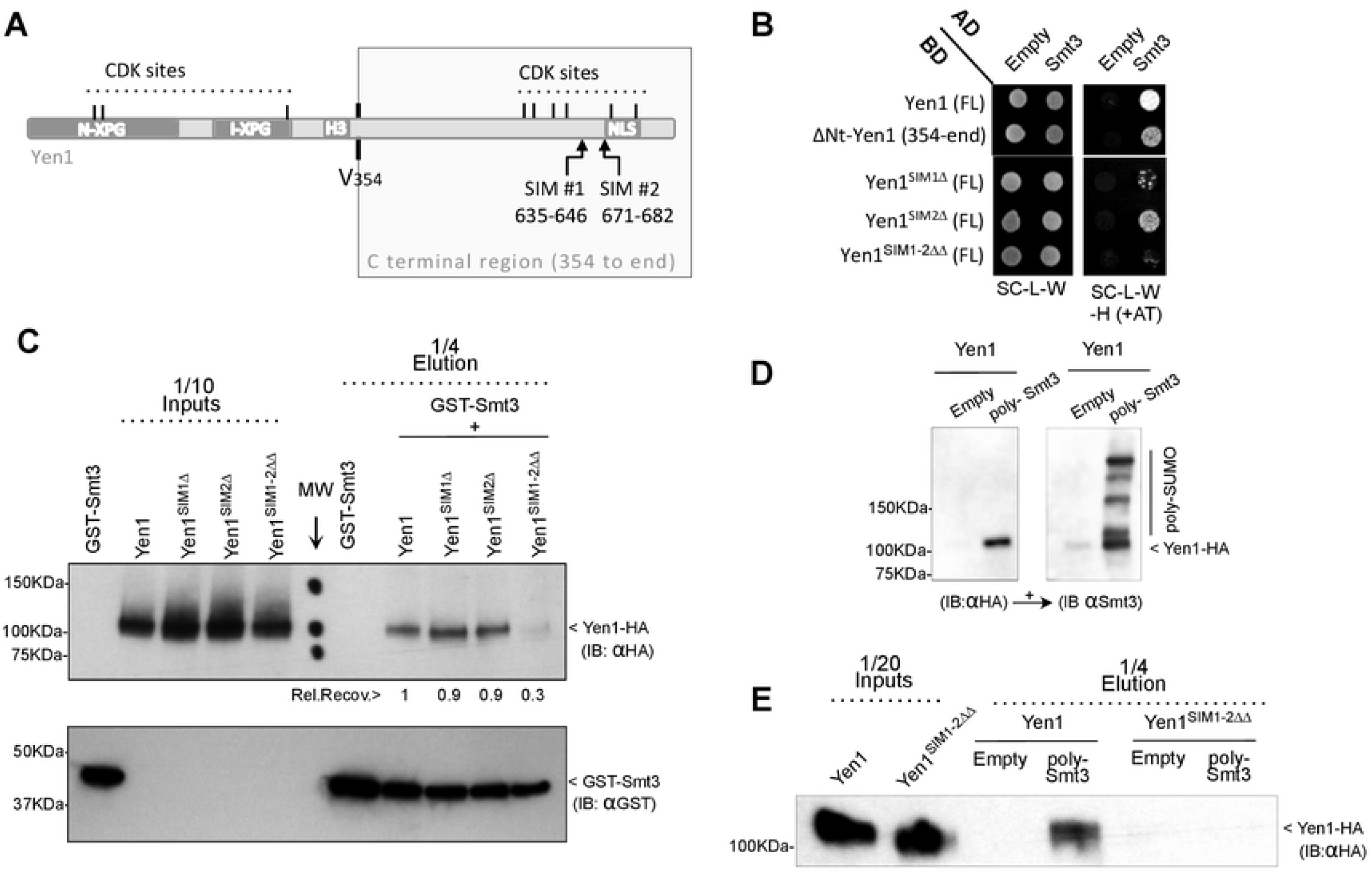
Yen1 contains two Sumo Interacting Motifs (SIMs) in its C-terminal domain. **(A)** Diagram showing the conserved domains of Yen1 and the positions of the regulatory Cdk1-phosphorylation sites. Amino acid 354 shows the cut-off point for truncated forms of Yen1 in Two-Hybrid assays. The two identified candidate SIMs are shown near the Nuclear localization sequence (NLS). **(B)** A Two-hybrid assay was performed with strains carrying the indicated Activator Domain (AD) and DNA Binding Domain (BD) fusions to test interaction between Yen1 and Smt3 (SUMO), and the Yen1 critical domains for such interaction. Mutations D635A D636A D637A for SIM1Δ and V675A E677A for SIM2Δ were introduced to test for the putative SIMs. SIM1-2ΔΔ is used for the combined mutations. Strains were grown on selective media lacking Leucine and Tryptophan and spotted in selective media also lacking Histidine to reveal interaction of the proteins fused to the AD and BD domains. Non-specific interactions were minimized by the addition of 3-aminotriazole (AT). **(C)** Purified GST-Smt3 was bound to a Glutathione resin and either purified wild-type or the SIM mutant Yen1 proteins were then loaded to the resin. The retained fractions of Yen1 were eluted after several washes and detected by immuno-blotting. **(D)** SUMO-retention assay using *in vitro* generated poly-Smt3 immobilized into a Cobalt HisPur Superflow agarose matrix. Purified Yen1 was added to the pre-bound Smt3 and after a binding-time and washing, columns were eluted in denaturing conditions and the eluates inspected by western blot for the presence of Yen1 (anti-HA, left panel) and the pre-bound Smt3 chains (right panel). **(E)** Immuno-blotting of the inputs and eluates of the retention assay (as in D) comparing the ability of wild-type or SIM-mutant Yen1 to bind poly-Smt3.

While mutation on the consensus SIM sites abolishes the interaction in the two-hybrid assay, this assay was suggested to reflect a covalent modification of Yen1 (40) and thus the effect detected in this test might be an indirect effect reflecting the loss of direct sumoylation of Yen1 related to the absence of non-covalent interaction of Yen1 to SUMO through its SIMs. Moreover, a possible SIM-mediated interaction would be difficult to be confirmed in this test if polymerization of Smt3 is needed or a Sumo covalent modification of a bridging partner is required for the growth read-out of the test. To better confirm the nature of the interaction lost in our mutants and thus validate the SIM sites, we used a pull-down approach (36). GST-Smt3 was over-expressed and purified from bacteria, and bound to a Glutathione resin. Purified Yen1 or its mutant SIM variants were then allowed to bind to the pre-bound GST-Smt3, and after several washes, the column content was eluted in denaturing conditions and inspected by western blotting (Figure 1C). Yen1 was detected in the eluates thus confirming its ability to interact non-covalently to Smt3, but was much less retained (30% compared to wild-type) when bearing mutations in both of its SIMs, while inactivation of only one of the two motifs did not significantly altered retention suggesting that at least *in vitro*, the presence of one single SIM at the Yen1 C-terminal region confers it ability to bind Smt3 (Figure 1C). While the GST-Smt3 column may indicate the ability of Yen1 to bind to monomers of Smt3 through its SIMs, the packed dispositions of GST-Smt3 in the column can mimick a poly-Sumoylated chain and contact Yen1 simultaneously in multiple SIM sites. To further evaluate the non-covalent binding of Yen1 to poly-Smt3 we decided to generate an affinity column containing poly-Sumo chains to test whether they have the ability to non-covalently bind to Yen1 and thus capture the protein. We generated Smt3 chains by adding Aos1-Uba2 and Ubc9 in a reaction with purified 6xHis-Smt3 (13), the resulting poly-Sumo chains were dialyzed and used to bind to a Cobalt HisPur Superflow agarose matrix, the poly-Smt3 coated matrix was used to test retention of Yen1-1xHA (Figure 1D). While the column retained the wild-type Yen1, the recovery of the protein mutated in both SIMs (Yen1^SIM1-2ΔΔ^) was greatly decreased (Figure 1E), thus confirming that Yen1 binds non-covalently to poly-Sumo chains, a binding that depends on the presence of the two identified Yen1’s SIMs.

### Strains carrying SIM-defective variants of Yen1 display increased sensitivity to DNA damage

We next aimed to understand the effect of the mutations in the SIM motifs on the ability of Yen1 to be normally regulated by Cdk1/Cdc14 and shuttled timely to the nucleus. Given the proximity of Yen1’s SIMs to its NLS, a C-terminal GFP fusion of the Yen1 mutants was monitored to see if any gross defect occurred for its nuclear shuttling (Figure 2). Both single and double SIM mutants presented nuclear exclusion in S-phase as the wild-type and were nuclear in late mitosis and G1 and the relative distribution of GFP intensity detected in the nucleus or in the cytoplasm at the different cell cycle phases was not changed between the different Yen1 variants (Figure 2B, Figure S2). We also analyzed the pattern of cyclic phosphorylation by synchronizing cells in G1 and analysing the mobility of Yen1 at different time points after its release (Figure 2C, Figure S2A). All the mutants displayed a normal cycle of phosphorylation in S-phase followed by gradual de-phosphorylation with only slight variations in the total amount of the protein all across cell-cycle phases. Next we asked whether the presence of an endogenous copy of the SIM mutants would compromise the ability of Yen1 to back-up for the functions of Mus81-Mms4 (5). The mutants were introduced into a *mus81*Δ background and tested for its sensitivity to an array of DNA damaging agents (Figure 2D and E). Cells with a double mutation *mus81*Δ *yen1*Δ are extremely sensitive to MMS at low doses, and they also present a moderate sensitivity to the radiomimetic drug Zeocin and to the replication stalling drug Hydroxyurea (HU) (13), both a mutation in the first SIM and the double SIM mutant significantly increased the sensitivity of a *mus81*Δ strain to MMS, while increased sensitivity to MMS after mutation in the second SIM alone was not significant (Figure 2D, 2E, Table S3). Mutation in both SIMs was necessary to see a moderate increase in sensitivity to Zeocin (Figure 2D, 2E). Mutation in both SIMs was also necessary to sensitize cells at 20mM HU and while individual SIM mutants sensitized cells to 40 mM HU, at such dose survival was already compromised by a two log difference in cells bearing both SIM mutations or lacking *YEN1* (Figure 2E). Yen1 has also been described to be essential in cells with a deletion on the *DNA2* helicase-nuclease in the presence of its suppressor *pif1*Δ (41-44). Similarly, cells carrying the *dna2-2* helicase-deficient allele rely on the activity of *YEN1* to process DNA intermediates in this background (41). Similar to what was already observed in *dna2-2* cells in the W303 background (41) we found that a *dna2-2* mutation is partially unviable with *yen1*Δ (Figure S3). The *dna2*-2 cells, while viable, present a strong heterogeneity of phenotypes likely associated to spontaneous accumulation of suppressors, as described (41) (Figure S3). Nonetheless, the introduction of the more severe of the SIM alleles, carrying simultaneous mutations in both SIMs, was viable in a *dna2-2* background, suggesting a minor role of non-covalent Sumo binding for the Yen1’s functions required in a *dna2-2* context. In agreement to the observation that *dna2*Δ *pif1*Δ strains do not significantly accumulate more Sumoylated Yen1 compared to the wild-type strain (13), we found that in a *dna2*Δ *pif1*Δ background the SIM defective allele does not increase the already strong sensitivity of cells to DNA damaging agents like MMS or HU (Figure S3).

**Figure 2.**
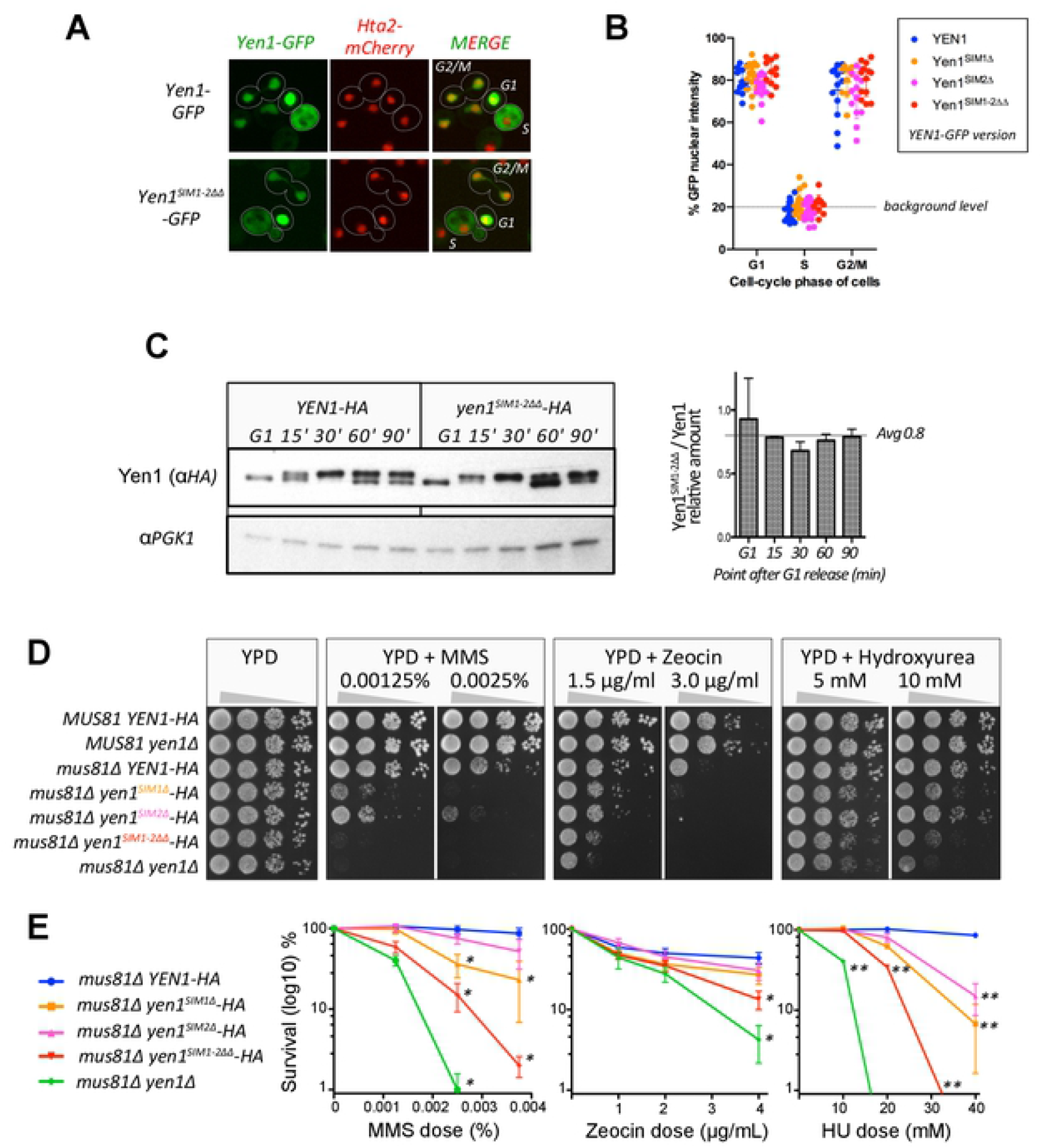
Mutation in Yen1 SIMs has no impact to its CDK1 regulation and nuclear shuttling but sensitizes cells to DNA damage. **(A)** Cells carrying an endogenous histone Hta2-mCherry marker and chromosomally –HA tagged versions of Yen1 wild-type and the different SIM mutants were transformed with a plasmid carrying an equivalent version of Yen1 fused with GFP at its C-terminal region. Cells were grown on selective media and observed using a spinning-disk microscope after a brief induction with galactose. Shuttling of the protein from cytoplasm to the nucleus can be observed in representative fields displaying cells with nuclear excluded Yen1 (S-phase and early G2-M) and nuclear localized Yen1 (anaphase to G1) for the indicated GFP-tagged proteins. **(B)** Quantification of the relative amount of GFP signal detected into the nucleus over the overall signal for the indicated strains in cells classified by their cell-cycle status **(C)** Strains with a chromosomally inserted copy of –HA tagged wild-type Yen1 or its double SIM mutant (Yen1^SIM1-2ΔΔ^) were synchronized with alpha factor and released into fresh medium to monitor the modification of the protein through the cell cycle by immunoblot (left). Both unmodified and phosphorylated Yen1 are indicated. Average levels of endogenous Yen1 were normalized with PGK1 protein in triplicate experiments (right). **(D)** Sensitivity to different DNA damaging agents and drugs was determined by spotting serial dilutions of strains carrying different Yen1 mutants in its SIM in a *MUS81* deleted background for the indicated media. **(E)** Survival curves to the agents tested in (C) were established by counting colony forming units of the different strains after plating in YPD containing the indicated doses of drugs in replicate trials. Survival was normalized per trial with its respective control YPD counts and the average % survival is plotted in the graphs (+/-SEM). Significance was estimated by the student T-test at P<0.05 (*) and P<0.01 (**), see Table S3. Additional data related to this figure is found in Figure S2.

### Mutation of the SIMs induces a sumo-less Yen1 phenotype *in vivo*

In other Sumoylated DNA repair proteins containing functional SIMs, the mutation of these motifs has an impact in the ability of the protein to be directly Sumoylated (36, 37). The results in the two-hybrid experiments suggested such an effect for Yen1 (Figure 1), which we have previously characterized to be Sumoylated ina Siz1/Siz2-dependent manner (13). To further confirm the absence of covalent sumoylation after impairment of Yen1’s SIMs we compared the Sumoylation levels of the wild-type and the SIM-defective Yen1 mutants by performing denaturing pull-downs of His-tagged Smt3 (Figure 3A). Yen1 Sumoylation peaks when cells are exposed to high MMS doses (13) and we reproduced Yen1 Sumoylation in these conditions for the wild-type protein (Figure 3A). Nonetheless, the fraction of Sumoylated Yen1 in the mutant in either the first SIM motif or the double mutant in the two SIM motifs was greatly reduced in conditions with similar input levels to 5% and 1% of the wild-type levels respectively (Figure 3A), mutation of SIM2 had a milder effect reducing the recovery of Sumoylated forms to ≈20% of the forms recovered in the wild-type (Figure 3A). The gradual effects detected for individual or combined mutations of both SIMs points to a concerted action of both motifs to promote Yen1 Sumoylation by allowing Yen1 non-covalent binding to Smt3. Despite its sensitivity to very low doses of MMS, cells carrying *dna2-2* did not spontaneously increase the yield of Sumoylated forms of Yen1 while we detected in such conditions under spontaneous damage a minor yield of Sumoylation in the presence of mutations on both Yen1 SIMs in either a wild-type or a *dna2-2* background (Figure S3).

**Figure 3.**
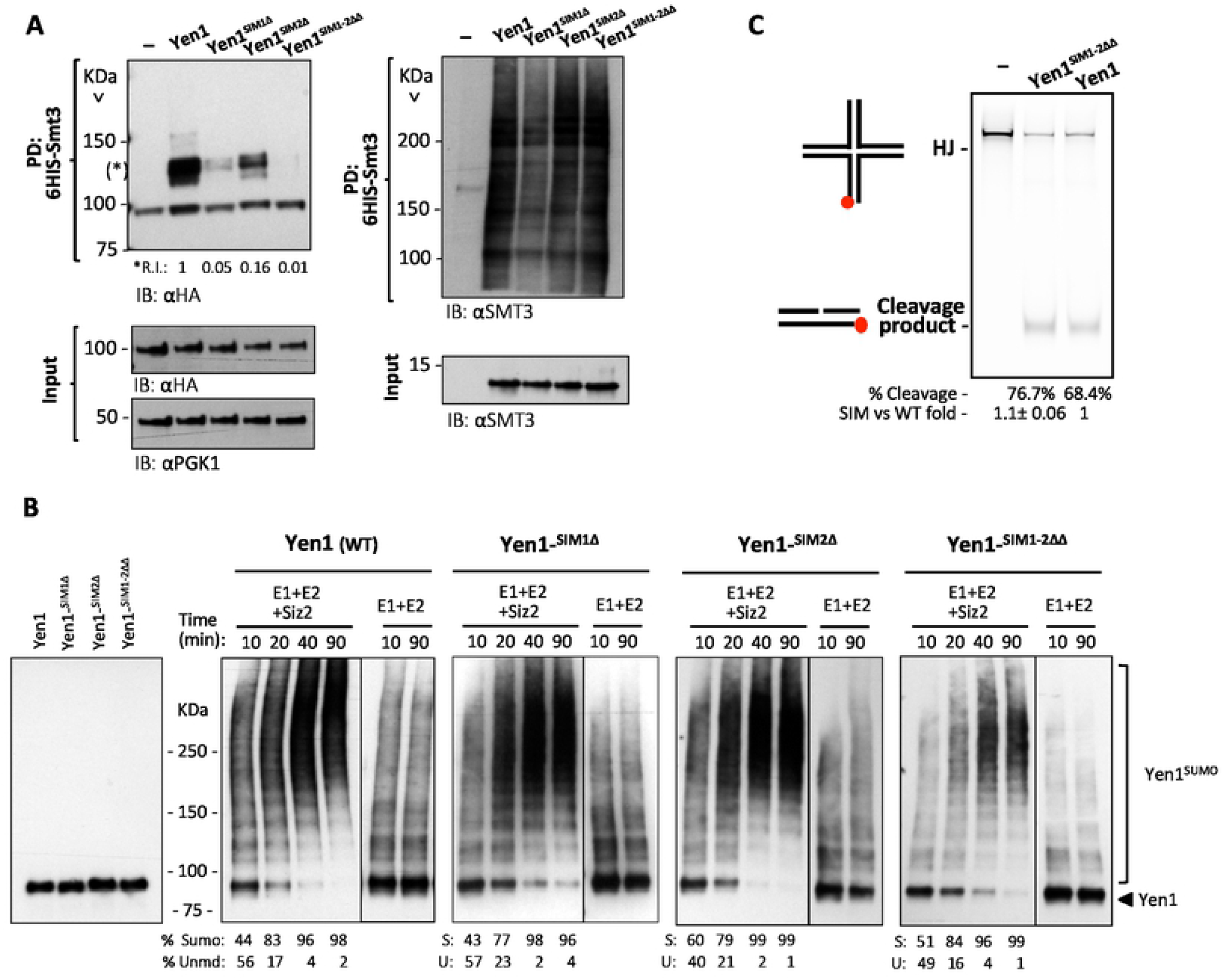
Mutation in Yen1’s SIMs does not alter its activity or its sumoylation *in vitro* but prevents sumoylation *in vivo*. **(A)** Strains carrying endogenous copies of –HA tagged wild type Yen1, Yen1^SIM1Δ^, Yen1^SIM2Δ^ and Yen1^SIM1-2ΔΔ^ mutants, with (+) or without (-) the plasmid pCUP-6xHIS-Smt3, were grown in the presence of MMS 0.3%. Cells were lysed and lysates subjected to a denaturing Ni-NTA pull-down followed by immunoblot analysis. Yen1 was detected by anti-HA. Membranes were subsequently probed with anti-Smt3. Prior to Ni-NTA pull-down, input samples were taken from the lysates and were analysed by immunoblotting for the levels of Smt3 induction (Anti-Smt3) and relative protein amounts (Anti-PGK1, Anti-HA) of each lysate (input panels). **(B)** Purified Yen1-HA and Yen1 SIM-mutant variants were subjected to an *in vitro* sumoylation reaction containing Siz2, Aos1-Uba2, Ubc9 and Smt3-3KR and ATP and subjected to Tris-Acetate PAGE for comparison of their sumoylation patterns after immunoblotting with anti-HA. The reaction without the ligase Siz2 is shown at the right of each panel. Quantification of the relative % of sumoylation forms (R.I.) and unmodified forms is shown at the bottom and more in detail in Figure S4 **(C)** Activity of Immuno-precipitated Yen1 was tested in a cleavage reaction using synthetic Holliday Junctions (HJ) made with oligonucleotides and labeled with Cy5. The DNA products were run in non-denaturing PAGE and revealed by the fluorescence of the Cy5 labeled oligonucleotide.

The mutations introduced to inactivate the SIM sites do not contain any lysine substitution, and SIM1 is not directly flanked by lysines in the immediate vicinity. To further confirm that the lack of Sumoylation was not due to un-adverted absence or un-accessibility of Sumoylation-target lysines in Yen1, we decided to test the mutant proteins in an *in vitro* Sumoylation reaction. After a reaction of the SIM mutants of Yen1 with Aos1-Uba2 and the conjugating enzyme Ubc9, a normal Sumoylation pattern was detected with the same ladder of bands of increasing sizes for all Yen1 variants (Figure 3B). We also performed a complete ligation reaction containing Siz2 as E3, increasing the yield of the reaction. Comparing the ligation reactions over time we couldn’t detect significant differences in the amount or timing of accumulation of the Sumoylated forms (Figure 3B, Figure S4) that in all the Yen1 variants achieved complete Sumoylation of the substrates at similar time points. We conclude that the presence of the SIM mutations does not preclude modification of any of the Yen1’s lysines targeted by the Sumoylation machinery.

The c-terminal domain of Yen1 containing the two SIMs is dispensable for a complete nuclease activity (45). Nonetheless, we verified that the mutation of both SIMs does not impair Yen1’s nuclease activity *in vitro* by using a synthetic Holliday Junction (46) as a substrate. Immuno-precipitated Yen1 was added to cleavage reactions, and we compared the yield of HJ cutting for either the wild-type Yen1 and the SIM defective mutant Yen1^SIM1-2ΔΔ^. The nuclease activity was undistinguishable for both the wild-type and the mutant, that were able to linearize the HJ substrate at similar rates (Figure 3C). Alteration of the SIM motifs at the C-terminal part of the protein seems thus not to alter the cutting efficiency of Yen1, whose nuclease and conserved XPG domains are present at the N-terminal part of the protein (Figure 1A). Nonetheless, the test did not take into account the disposition of the junction in a chromatin context in the cell that could influence the ability to cut HJ *in vivo*.

### Localization to spontaneous and induced sites of activity is impaired by inactivation of the Yen1’s SIMs

Sumoylation and interaction with SIMs has been proposed as a way to enforce a cascade of interactions to foster recruitment of factors to specific subcellular locations (35). We decided to determine if the impairment of the Yen1 SIMs was altering its ability to cluster in foci that are observed to occur either spontaneously or induced by DNA damage (4, 13). C-terminal GFP tagged versions of Yen1 were compared for its foci distribution (Figure 4, Figure S5, Table S4 and S5), the number of cells showing spontaneous foci in a wild-type strain ranges around 10%, while only 3% of the cells in the double SIM mutant displayed foci in strains with a functional Mus81-Mms4 (Figure 4B). The effect was more marked when observing the spontaneous foci in a *mus81*Δ strain, where cells displaying foci decreased from 30% to 3-4% in the SIM mutant (Figure 4B). These foci were equally decreased in the presence of exogenous damages, suggesting that Yen1 SIMs are equally important to properly localize the protein to spontaneous damaged sites and exogenous damages sites (Figure 4B, Table S6 and S7).

**Figure 4.**
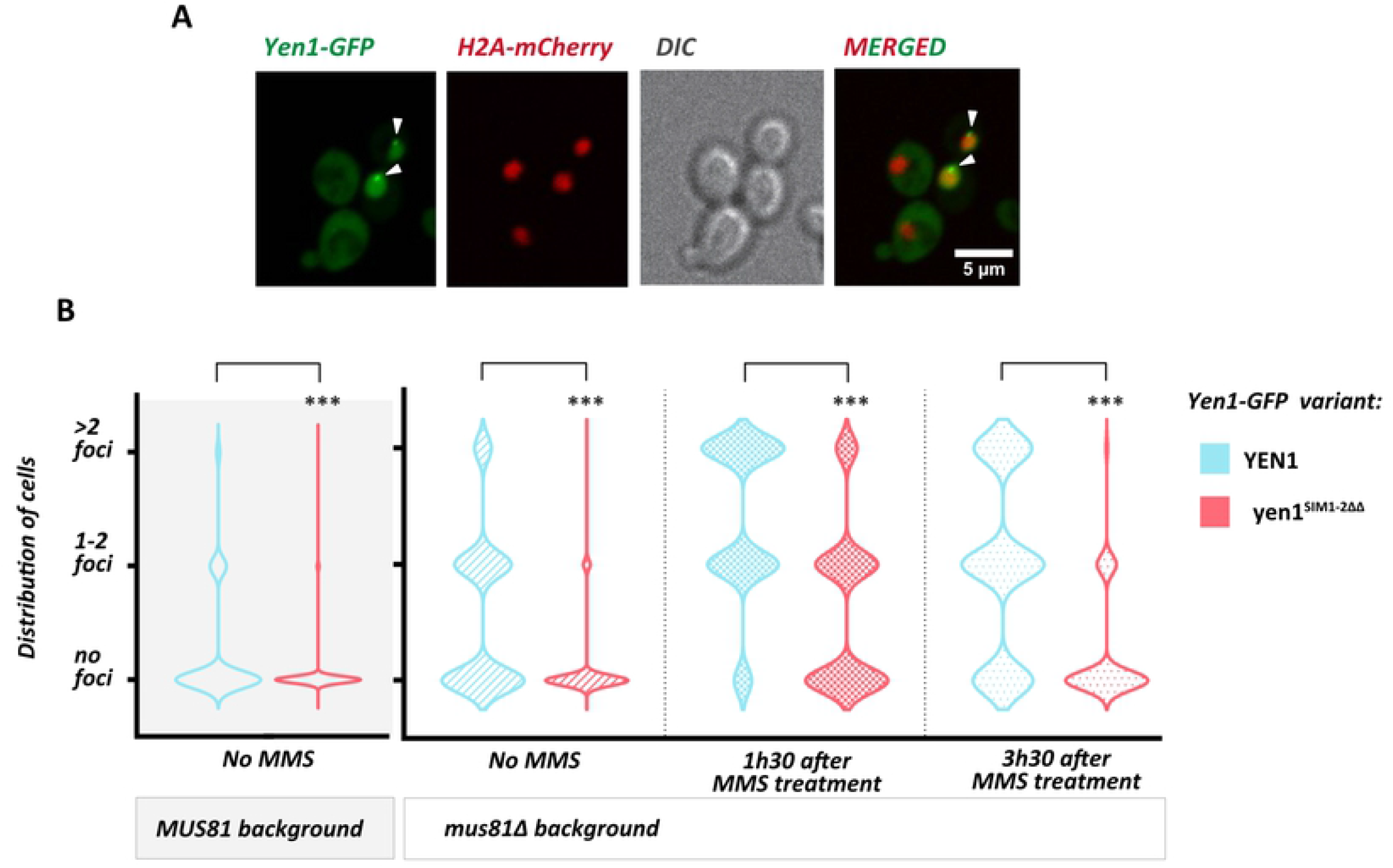
Mutation in the SIMs of Yen1 prevents foci accumulation in G2/M. **(A)** Cells with an endogenous copy of Hta2-mCherry and YEN1-HA expressing Yen1-GFP from an inducible vector were observed under a spinning-disk microscope after a brief induction with galactose. The white triangles denote the presence of Yen1-GFP foci. **(B)** Chromosomally tagged Yen1-HA wild-type and SIM mutant in the indicated genetic backgrounds and carrying its corresponding Yen1-GFP expressing plasmid (blue graphs for WT copy, red for SIM mutant) were observed under the microscope after a brief induction. Cells from the indicated conditions were classified according to its cell cycle phase and the presence or absence of Yen1 foci. Violin Plots display the distribution of G2/M cells showing no foci, 1-2 foci or more than 2 foci for each strain. Counting was performed for over 400 distinct G2/M cells for each strain over several independent trials. Asterisks represent statistical significance in a χ2 test p<0.0001 (***), see tables S4, S5, S6 and S7 for details.

### Absence of Yen1’s SIMs prevent accumulation of the Yen1 fraction targeted by Slx5-Slx8

We have demonstrated in a previous work a role for the Slx5-Slx8 Sumo-targeted ubiquitin ligase in the removal of a subset of Yen1 from the nucleus during the transition from G1 to S phase (13). As a result, cells defective in Slx8 show a persistence of Yen1 foci, which are detected in large numbers even in the presence of functional Mus81-Mms4 (13). We wondered if the absence of proper localization in the SIM mutant would prevent Yen1 accumulation in the absence of Slx5-Slx8. According to our expectations, a deletion of *slx8*Δ in the strain bearing the mutations in the Yen1’s SIMs did not increase the number of Yen1 foci, thus suggesting mis-localization of Yen1 in the mutant strain prevents the need for Slx5-Slx8 targeted removal (Figure 5A, Table S8). The persistence of the Slx5-Slx8 targeted Yen1 fraction can also be detected by performing a cycloheximide chase during a synchronous release from G1, and observing the degradation of Yen1 under inhibition of *de novo* protein synthesis in this cell-cycle interval (13). In a *slx8*Δ background, the double SIM mutant protein was degraded during such cycloheximide chase faster than the wild-type protein (Figure 5B), indicating that the lack of accumulation of Yen1 at nuclear sites does also dispense for a targeted removal of this subset of the nuclease, that is removed timely even in the absence of Sumo-targeted ubiquitination by Slx5-Slx8.

**Figure 5.**
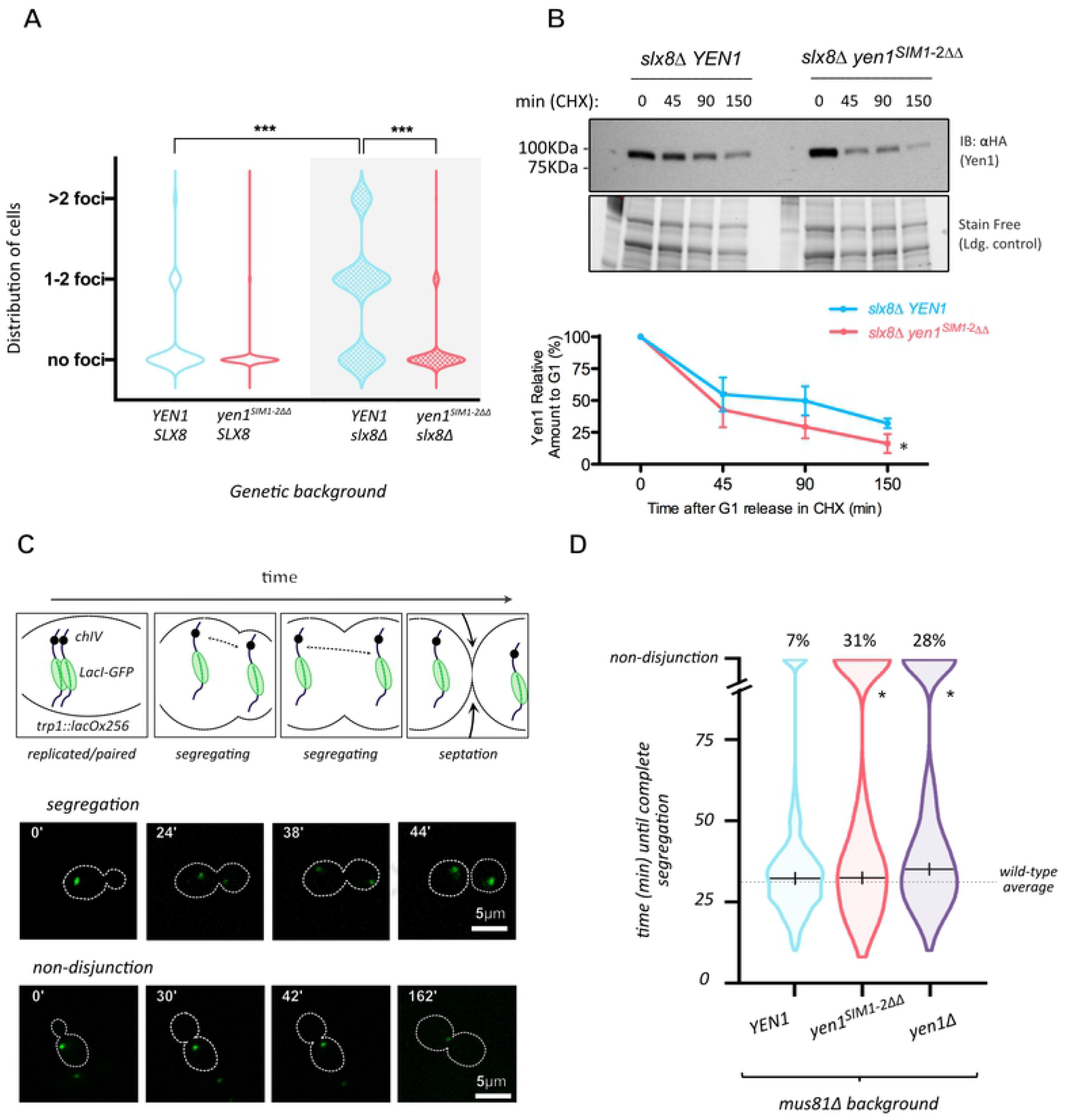
Mutation in the SIMs of Yen1 prevents foci accumulation in *slx8*Δ cells during G2/M and impacts chromosome segregation in a *mus81*Δ background. **(A)** Cells containing a deletion on *SLX8* were observed for their distribution of foci of the different variants of Yen1-GFP. Violin plots display the distribution of cells and asterisks denote statistical significance in a Chi-square test at P <0.0001 (***), see table S8. **(B)** Cells from the indicated genotypes were arrested in G1 and released in the presence of cycloheximide with samples being taken at the indicated time points. Total protein extracts were inspected by immunoblot for the presence of Yen1-HA and their intensity quantified relative to the loading control obtained by stain-free imaging of the gels (BioRad). Relative amounts of Yen1 are plotted in the graph to facilitate comparison (+/-SD). **(C)** Diagram showing chromosome segregation in cells harboring a lacO/GFP-LacI array tag on chromosome VII. To discriminate cells with timely chromosome segregation from those presenting aberrant segregation (delayed segregation or non-disjunction) a 2 h limit of observation was implemented. Two sets of representative actual images of a normal segregation pattern and a non-disjunction pattern are shown below the diagram. **(D)** Over 400 cells per strain were individually counted and are represented in violin plots according to the time spent to segregate the lacO/lacI array. The median segregation time is indicated excluding cells with non-disjunctions. Statistical relevance of the differences observed between the number of non-disjunctions of the different strains was determined by the Chi-square test at P<0.0001, see table S9.

### Unpaired Sumo-directed localization induces an increase in untimely chromosome segregation

The presence of both *mus81*Δ and *yen1*Δ deletions makes cells synergistically sensitive to drugs like MMS, as stated before, and also increase their spontaneous number of chromosome mis-segregations monitored either by dedicated genetic systems (5, 47) or by a direct observation of fluorescent-tagged chromosomes during mitotic divisions (13) (Figure 5C). We compared the mitoses of single *mus81*Δ cells to that of cells carrying *mus81*Δ and the allele with the two mutated SIMs. Similar to what we could observe in a *mus81*Δ *yen1*Δ control strain, *mus81*Δ cells with the mutated Yen1 SIMs displayed an increased number of segregation issues. As it can be observed in the violin plots displaying the time that individual monitored cells spent to fully segregate the fluorescent tag, the Yen1 double SIM mutant does segregate chromosomes in a similar average time than the wild type for those cells completing full segregation (Figure 5D) but about 30% of the cells carrying these Yen1 SIM mutations in the *mus81*Δ background were unable to resolve its segregation within the time of video-microscopy observation (2 h), and were classified as non-disjunctions (Figure 5D, Table S9). The high number of chromosome mis-segregation detected for the Yen1 mutant variant in a *mus81*Δ background is in line with that observed in cells completely lacking Yen1 clearly pointing to a faulty function of Yen1 when SIMs are absent.

### Mutation in the SIMs of Yen1 reduces the formation of crossing-over after a single DSB

To further browse the implications of the presence of a defective Sumo-interacting Yen1 allele for the actual resolution of recombination intermediates, we decided to analyse the level of crossing-over (CO) formation in two widely used tests that estimate the CO levels after a single DSB formation (3, 5, 48) (Figure 6A and D). In accordance with the increased sensitivity to different DNA damaging agents observed for the Yen1 allele carrying the SIM mutations when combined with a *mus81*Δ background (Figure 2), we detected a decreased formation of crossovers in this genotype after the induction of a single DSB in a diploid tester strain (5), half-way to the phenotype observed with a double mutant carrying both deletions in *mus81*Δ and *yen1*Δ (Figure 6B and C, Table S10). The decrease in crossover formation was paralleled by an increase in Break-Induced Replication (BIR) events (Figure 6B and C, Table S10). Using an ectopic recombination assay (48) (Figure 6D and E), we detected a decrease in viability after the induction of an HO cut site in chromosome II in the *mus81*Δ *yen1*^*SIM1*-2ΔΔ^ strain, already signalling a defective crossover resolution resulting in a number of unviable events (Figure 6F). This survival decrease probably reflects a BIR increase that in this test leads to lethality by the loss of essential genes in the chromosome II distal arm. The number of crossovers quantified by southern blotting analysis of the survivors showed a nearly 50% reduction in the crossover yields in *mus81*Δ *yen1*^*SIM1*-2ΔΔ^ cells (Figure 6E and F, Table S11), not significantly different from the levels detected in *mus81*Δ *yen1*Δ cells.

**Figure 6.**
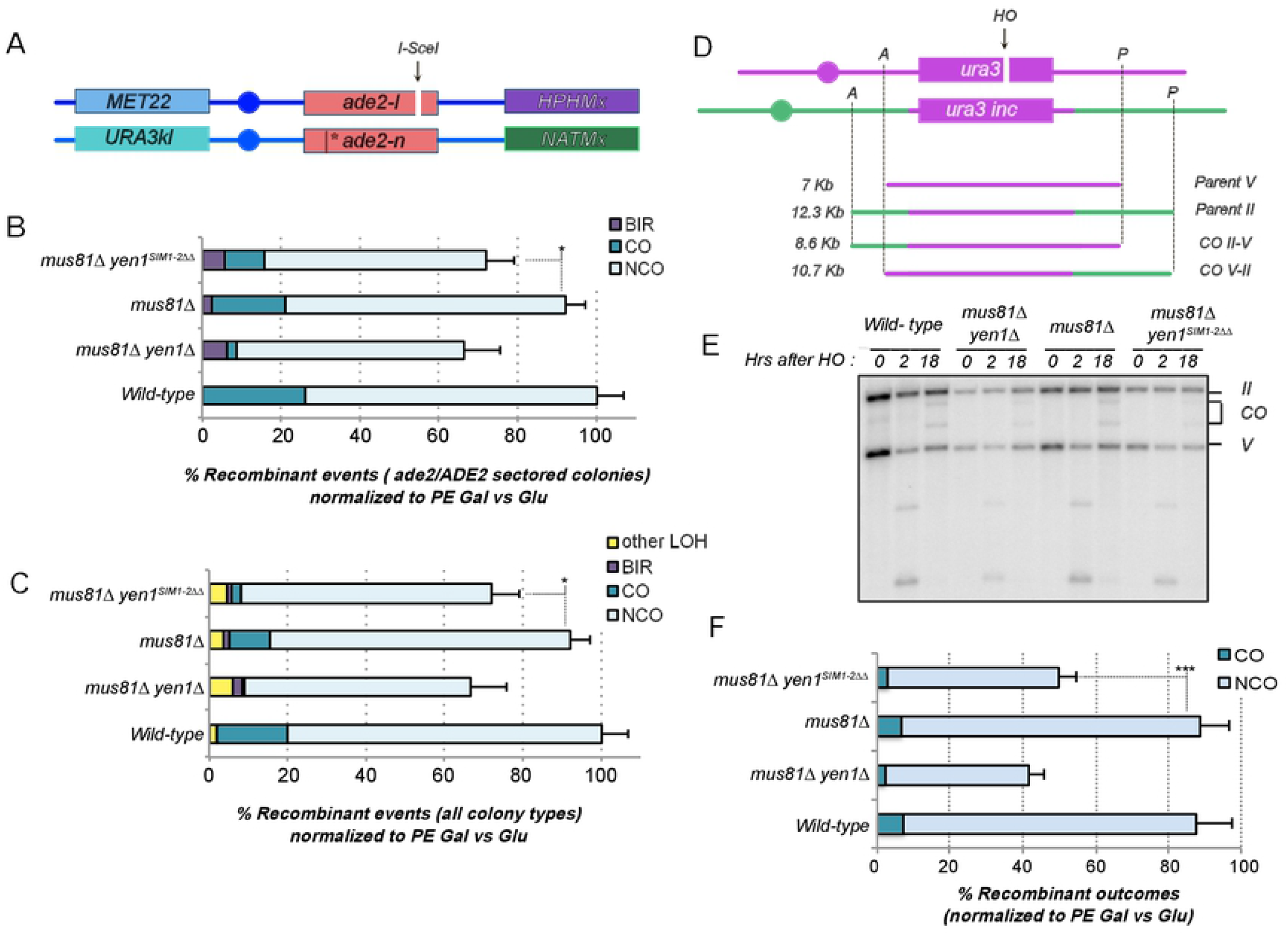
Crossover formation is impaired in cells containing the mutant version of Yen1 inactivating both SIMs in a *mus81*Δ background. **(A)** Diagram showing the chromosome XV based DSB-induced recombination reporter. **(B)** Recombination outcomes in red-white (ade2/ADE2) sectored colonies of the indicated strains, normalized to their Plating Efficiency (PE) in Galactose compared to Glucose. **(C)** Recombination outcomes combining the results obtained for all types of colonies (full red, full white and sectored) of the indicated strains, normalized to their Plating Efficiency (PE) in Galactose compared to Glucose. Statistical significance for B and C was determined by the Chi-square test at P<0.05, see table S10. **(D)** Diagram showing the chromosome II-V based ectopic DSB-induced recombination reporter and its expected outcomes during physical analysis. **(E)** Representative southern blot analysis of the indicated strains after genomic DNA digestion with the restriction enzymes as highlighted in diagram D and hybridization with a radiolabeled probe against the *URA3* locus. **(F)** Quantification of at least three independent southern blot analyses is plotted relative to PE (Galactose vs Glucose). Statistical significance was determined by the Student T-test at P<0.05, see table S11.

## Discussion

In the present work we aimed to understand whether the Yen1 nuclease depends on interactions with Sumoylated partners to be able to act accurately and promptly on its substrates. We have demonstrated that in addition to being Sumoylated, Yen1 is also able to interact non-covalently to Sumoylated chains and Sumo monomers through at least two Sumo-interacting motifs in its C-terminal region (Figure 1). Previous reports suggested covalent Sumoylation occurring in the C-terminal region of Yen1 was responsible for the Smt3-interaction detected in a two-hybrid assay (40), our results are in agreement with the original observation of a C-terminal motif mediating such Smt3-interaction but we conclude this results are compatible with a non-covalent interaction mediated by the two SIMs, leading to a covalent modification of Yen1. We have demonstrated such non-covalent interaction with dedicated retention assays using either immobilized GST-Smt3 or pre-polymerized poly-(6HIS)-Smt3 coupled to a Cobalt HisPur Superflow agarose matrix (Figure 1) in a strategy similar to that used to validate other sumoylated protein’s SIMs (36, 49-51). Moreover, we further confirmed the nearly complete loss of *in vivo* Sumo covalent modification in the SIM defective Yen1 using 6His-Smt3 denaturing pull-downs (Figure 3A). Our Yen1 SIM mutant acts thus as an *in vivo* Sumo-less variant without requiring a large number of Lysine substitutions, which can sometimes result in the protein’s destabilization. While direct Yen1 sumoylation depends strongly in the presence of the identified SIM motifs, we conclude those are not required to mediate interaction to the sumoylation machinery *per se*, as we can observe full sumoylation patterns after an *in vitro* sumoylation reaction (Figure 3B) and we also detect residual forms of sumoylated Yen1 in our denaturing pull-down assays (Figure 3A) displaying the regular band pattern of *in vivo* sumoylated Yen1, and not the absence of these forms that is obtained when Siz1 and Siz2 are removed (13). However, we conclude that this direct sumoylation of Yen1 is largely prevented in the cells by a faulty localization *via* SIMs to specific nuclear sites as suggested by the inability of the SIM mutant to accumulate in foci in sub-nuclear localizations previously characterized (13) (Figure 4). Accordingly, while the absence of the SIMs in Yen1 has no effect on its catalytic activity (Figure 3C), we have detected a sub-optimal function of these mutants in the cells, leading to phenotypes of chromosome mis-segregation and DNA damage sensitivity similar to those observed for a null mutant in combination with a deletion of the partially redundant cell’s major resolvase activity mediated by the heterodimer Mus81-Mms4 (Figures 2, 4 and 5) (5). The impaired localization not only correlates with a sub-optimal function of Yen1 in response to spontaneous damages under normal growth conditions and exogenous treatments with genotoxic agents, but also decreases the number of crossing-over that can be observed after a single DSB induction in two different settings (Figure 6). While inactivation of both SIMs identified in the C-terminal region is necessary to impair non-covalent binding to Sumo, the mutation we introduced in SIM1 seems to achieve a stronger phenotype alone than the one in SIM2. Nonetheless, we did not perform an optimized serial mutation of each motif and thus we can’t exclude that both SIM motifs contribute with equal importance to Sumo binding in the cells.

Our results are in line with group modification (35) and would suggest a local enrichment of multiple Sumoylated proteins together with free Smt3 and the Sumoylation machinery when persistent recombination intermediates are revealed during anaphase. While Sumoylation has been previously shown to play important roles in the fine-tuning of DNA repair processes, our study highlights the importance of Sumoylation for genome maintenance processes occurring in anaphase, and probably disconnected to previous Sumoylation cascades influencing HR proteins. Several key proteins acting in anaphase, like Condensin subunits and chromosomal passenger complex (CPC) components, have been described to be Sumoylated (20, 52, 53). It is thus of great interest to continue studying Yen1 functional interactions in conditions that are greatly transient an ephemeral, and determine which other factors ensure prompt Yen1 recruitment to its activity sites during its anaphase activity window, thus influencing the delicate balance between chromosome segregation and genome integrity.

## Materials and Methods

### Yeast Strains and Growth Conditions

*S. cerevisiae* strains used in this study are derivatives of the W303c background and are listed in Table S1. The Yen1-FX-GFP allele was made by inserting a Factor X site and the GFP epitope from pGAD-Yen1-GFP(54) between amino acids D753 and S754 at the C-terminus of Yen1 using dedicated oligonucleotides and was cloned into TOPO-pYES2 (Invitrogen) to allow controlled expression by Galactose induction, all plasmid derivatives are listed in Table S2. Mutants in the different designated loci where either obtained by crossing or by gene replacement with the indicated selective cassettes. Cells were typically grown in YP (1% yeast extract; 2% peptone) or SC media with alternatively 2% glucose, 2% raffinose or 2% galactose in strains under inducible conditions. A modified medium (SC with 0.17% YNB without ammonium sulfate, 0.1% proline and 0.003% SDS) was used for the Smt3 pull-down assays.

### Western Blot analyses

If not stated otherwise, proteins were extracted by the TCA (Trichloroacetic acid) method. For routine monitoring, samples were loaded into 7.5% Tris-Glycine stain-free pre-casted gels (BioRad). Samples from pull-downs analyses were loaded into 3-8% gradient NuPAGE Tris-Acetate gels (ThermoFisher). Gels were transferred using a semi-dry transfer machine (BioRad) to PVDF membranes and hybridized with the appropriate antibodies in 5% w/v nonfat dry milk, 1X TBST buffer. Antibodies for anti-HA-HRP (3F10, Roche), anti-Smt3 (B. Palancade), anti-Pgk1-HRP (22C5D8, Abcam) were used at the suggested dilutions and revealed using an ECL reagent (Advansta). When required, HRP-conjugated secondary antibodies from Cell Signaling were used at 1/10000 dilution.

### Microscopy and Cell Biology Methods

Live cell imaging was performed with a Spinning Disk Confocal Microscope (CSU-W1, Yokogawa), with an electron multiplying charge device camera (ANDOR Zyla sCMOS) and a ×60/1.35 numerical aperture objective at 30 °C. Cells were centrifuged and plated as a droplet between an SC agarose pad and a glass slice (55). Images were recorded with 17 z-sections with 0.5 µm spacing for each wavelength at a time. Video recordings were built with images taken every 2 minutes. Metamorph was used for image acquisition, and analysis was performed using Image J-Fiji (56).

For Yen1 foci observations, cells were grown in SC medium without uracil (SC-URA 2% raffinose), GFP-Yen1 was induced in a short burst of 30 min with Galactose at 2%, followed by addition of Glucose at 2%. For acute exposure to DNA damage, cells were treated with MMS 0.01% for 15 min at room temperature and were washed once with fresh SC-URA 2% glucose before continuing the experiment. Aliquots were taken at the indicated times. Cells showing an accumulation of spots were measured at maximum projection of the GFP channel. Statistical significance was determined by the χ2 test using contingency tables with the number of cells observed in each different category.

For segregation monitoring using strains with the lacO/GFP-LacI array, all cells were recorded for a duration of 2 h minimum in their agarose pads. Individual cells were identified with an ongoing chromosome segregation. To determine segregation duration a start point was determined as the signature S-phase bud was the smallest yet discernable. At this point, only one foci of GFP-tagged chromosome fluorescent markers is visible. The cell is followed until the dot separates in two and resides durably in the daughter cells. The ending time point is taken at the last frame of definitive separation of the fluorescent foci. The duration of the movement of the two separate dots was reported for each individual cell under monitoring, cells with dots moving together for the whole duration of the time-lapse were assigned as non-disjunction and their segregation time was not used to establish the average segregation time.

### Sumoylation assays and Smt3-bound retention assay

In *ex vivo* Sumoylation assays, the wild-type or mutant Yen1-HA was produced from a pYES2 vector and immuno-precipitated from cell lysates as described (13). Eluates were subjected to Sumo conjugation and ligation as described (57).

For Smt3-retention assays, 6x-His-Smt3 was purified from BL21 *E*.*coli* cells using a Ni-NTA affinity column (Qiagen) following manufacturer indications. Smt3 protein was eluted with 250 mM imidazole before being dialyzed using a G2 Slide-a-Lyzer cassette (Thermo Fisher) with a 10 kDa cut-off. Purified Smt3 was subjected to a self-conjugation reaction by adding Aos1-Uba2 and Ubc9 and ATP as described (57) and the reaction was subjected to a second purification in a Cobalt HisPur Superflow agarose matrix (Thermo Fisher) to generate poly-Smt3 retention column. Equal amounts of Yen1 or its mutant were added to the non eluted matrix containing poly-Smt3 bound in E buffer (20 mM NaH_2_PO_4_, 300 mM NaCl, 5 mM Imidazole, pH 7.4) and binding was allowed for 60 min at 4°C. Mini-Columns were then centrifuged to remove the buffer and non-retained proteins, washed 5 times in washing buffer (E buffer 12.5 mM Imidazole) and eluted in denaturing conditions with Laemmli buffer at 95°C. The eluates were loaded into 4-15% SDS-PAGE gradient gels and immunoblotted. GST-Smt3 retention assays were performed as described (36) with purified GST-Smt3 obtained by expression of pGEX-4T1-Smt3 into BL21 *E*.*coli* cells. Briefly, purified GST-Smt3 was incubated with glutathione matrix during 40 min at 4°C in GST buffer (25 mM Tris-HCl (pH8.0), 150mM NaCl, 1mM DTT). Then matrix was washed before to load purified wild-type or mutated Yen1. Mixtures were incubated 3 min at 4°C before to wash the glutathione matrix and elute retained proteins for HA and anti-GST immuno-bolt analysis.

### Cycloheximide chase experiments

Cycloheximide chase experiments were essentially done as reported (13). Cultures grown in SC complete modified media (0.1% proline 0.017% YNB without ammonium sulfate, 0.0003% SDS) were diluted to OD_600_=0.2 and synchronized with alpha factor (3 µM) for 2 h. Once synchronized, cells were treated with cycloheximide (250 µg/ml) in fresh media, to inhibit proteins new synthesis, and released from the G1 arrest. Samples were taken at indicated time points and analyzed by TCA extraction and western blotting.

### Denaturing Histidine pull-downs

For 6xHIS-Smt3 pull-downs, strains containing the expression vectors or the control empty plasmid were grown in SC-LEU modified medium (0.1% proline, 0.017% YNB without ammonium sulfate). Cells were allowed to grow to OD_600_=0.3 when CuSO_4_ was added at 100 µM final concentration in a volume of 100 ml. After 1 h, MMS was added to 0.3% and cells were collected 3 h later. Cells where lysed under denaturing conditions and Sumo-conjugated proteins where isolated and analyzed by western blot using a Nu-PAGE Tris-acetate 3-8% gradient gel, basically as previously described (13). In *dna2-2* strains (LEU2), a plasmid expressing under Galactose induction Flag-6His-Smt3 (URA3) was used instead of the Cu-inducible.

### Synthetic DNA substrates and Yen1 resolvase activity assays

The synthetic HJ-X0 was prepared by annealing the Cy5-X0-1, X0-2, X0-3 and X0-4 oligonucleotides (Sigma-Aldrich) in a buffer containing 50 mM Tris-HCl (pH 7.5), 10 mM MgCl_2_, 100 mM NaCl as described (46). The annealing product was analyzed in a native PAGE to verify the presence of a HJ structure. To test Yen1 activity, an enzymatic reaction was performed in 10 µl cleavage buffer (50 mM Tris-HCl (pH 7.5), 1 mM MgCl_2_, 1 mM DTT) containing 25 nM of Cy5-labled HJ X0 substrate, and equal amounts of immunoprecipitated Yen1 or its SIM-defective mutant. After incubation at 30°C for 1 h, the reaction was stopped by adding 2.5 µl of stop buffer (100 mM Tris-HCl (7.5), 50 mM EDTA, 2.5% SDS, 10 mg/ml proteinase K) and further incubated for 30 min at 37°C. Cleavage products were migrated in 10% native PAGE, scanned using a Typhoon FLA 9500 Biomolecular Imager and the images were analyzed with ImageQuant (GE Healthcare).

### DSB-induced recombination assays

The diploid recombination assays were performed as described previously (for a detailed protocol see (47)). The reporter diploid strains that contain 2 *ade2* hetero-alleles were cleaved by induction of I-SceI in its *ade2*-I allele and allowed to repair with its ade2-n allele under non-selective conditions to give rise to either ADE2 or *ade2*-n repair products in three types of colonies (red, white and sectored). Each recombinant colony was scored with its two recombination events (for each repaired sister chromatid) and considering the possible segregation patterns in daughter cells. Frequencies of the recombination events were normalized to the galactose vs glucose plating efficiency.The distribution of CO and NCO in the ectopic recombination assay based in chromosomes V and II (48) were addressed by southern blot hybridization of ApaLI-PvuII digested genomic DNA from cell populations growing in YP-Raffinose after galactose induction of HO. Membranes were hybridized with a *URA3* radiolabeled probe and results were normalized relative to the galactose versus glucose plating efficiency of the strains as described (3).

## Acknowledgments

We thank L.S. Symington, J. Campbell, M. E. Budd, K. Dubrana and S. Marcand for generous gifts of yeast strains and plasmids. We thank B. Palancade and S. Brill for supplying reagents for the Sumoylation assays. We thank the members of the UMR9019 CNRS for the useful discussions and suggestions. We thank Fondation ARC pour la Recherche sur le Cancer and donators from Natixis for their support of our research. GM is a full-time INSERM researcher at the CNRS. IT is a full-time CNRS researcher. HD has benefited from a doctoral fellowship from the French Education Ministry.

## Data Availability

The data that support the findings of this study are available from the corresponding author upon reasonable request

